# PhyloFunc: Phylogeny-informed Functional Distance as a New Ecological Metric for Metaproteomic Data Analysis

**DOI:** 10.1101/2024.05.28.596184

**Authors:** Luman Wang, Caitlin M. A. Simopoulos, Joeselle M. Serrana, Zhibin Ning, Yutong Li, Boyan Sun, Jinhui Yuan, Daniel Figeys, Leyuan Li

## Abstract

**Background:** Beta-diversity is a fundamental ecological metric for exploring dissimilarities between microbial communities. On the functional dimension, metaproteomics data can be used to quantify beta-diversity to understand how microbial community functional profiles vary under different environmental conditions. Conventional approaches to metaproteomic functional beta-diversity often treat protein functions as independent features, ignoring the evolutionary relationships among microbial taxa from which different proteins originate. A more informative functional distance metric that incorporates evolutionary relatedness is needed to better understand microbiome functional dissimilarities.

**Results:** Here, we introduce PhyloFunc, a novel functional beta-diversity metric that incorporates microbiome phylogeny to inform on metaproteomic functional distance. Leveraging the phylogenetic framework of weighted UniFrac distance, PhyloFunc innovatively utilizes branch lengths to weigh between-sample functional distances for each taxon, rather than differences in taxonomic abundance as in weighted UniFrac. Proof-of-concept using a simulated toy dataset and a real dataset from mouse inoculated with a synthetic gut microbiome and fed different diets show that PhyloFunc successfully captured functional compensatory effects between phylogenetically related taxa. We further tested a third dataset of complex human gut microbiomes treated with five different drugs to compare PhyloFunc’s performance with other traditional distance methods. PCoA and machine learning-based classification algorithms revealed higher sensitivity of PhyloFunc in microbiome responses to paracetamol. We provide *PhyloFunc* as an open-source Python package (available at https://pydigger.com/pypi/PhyloFunc), enabling efficient calculation of functional beta-diversity distances between a pair of samples or the generation of a distance matrix for all samples within a dataset.

**Conclusions:** Unlike traditional approaches that consider metaproteomics features as independent and unrelated, PhyloFunc acknowledges the role of phylogenetic context in shaping the functional landscape in metaproteomes. In particular, we report that PhyloFunc accounts for the functional compensatory effect of taxonomically related species. Its effectiveness, ecological relevance, and enhanced sensitivity in distinguishing group variations are demonstrated through the specific applications presented in this study.

## Background

The human body is inhabited by trillions of microorganisms that collectively shape the functionality of our complex internal ecosystems, primarily through protein activity [1, 2]. The integration of taxonomic composition, functional activity, and ecological processes offers valuable insights into the dynamic responses of the microbiome [3]. Metaproteomics stands out among omics approaches by directly measuring protein expression, providing unparalleled insights into the functional activities of microbial communities [4]. Beta-diversity is a metric to measure the degree of dissimilarity between ecological communities [5], and it has been applied to metaproteomics datasets by assessing the variation in abundances of function or taxonomic composition inferred by protein biomass [6, 7]. Microbiome *functional beta-diversity* refers to the variation in functional gene/protein patterns between microbial communities across different environments or conditions [8, 9]. It has been used to gain valuable insights into the patterns and variations across different metaproteomes [10].

Beta-diversity metrics are typically calculated without incorporating phylogenetic information [11]. Commonly used beta-diversity measures, such as Bray-Curtis dissimilarity [12], Jaccard distance [13], Euclidean distance, etc., rely solely on the abundance or presence/absence of taxa or functions within communities. These metrics are useful for comparing compositional features across samples, but do not account for the phylogenetic relationships or evolutionary history of microbial taxa within the communities. In contrast, the UniFrac distance [14, 15] was specifically developed for microbiome composition data, considering both the abundance of taxa (or their presence/absence in the case of unweighted UniFrac), and their phylogenetic relatedness. This makes UniFrac more biologically meaningful in reflecting microbial community differences compared to methods that rely solely on taxa abundances [14, 15]. However, the UniFrac distance is measured by taxonomic presence/abundance and does not incorporate any functional information of the microbiome. Functional compensatory effect between phylogenetically related species describes a dynamic process where closely related species adjust their functional roles to maintain overall ecosystem functionality. Therefore, the inclusion of functional information can provide a more ecologically relevant perspective compared to relying solely on species abundances. More recently, computational algorithms Phylogenetic Robust Principal-Component Analysis (Phylo-RPCA) and Phylogenetic Organization of Metagenomic Signals (POMS) have been developed to integrate phylogenetic information with metagenomic functional profiles [16, 17]. Despite these advancements, these diversity metrics are derived without considering whether these genes are expressed or not. In other words, these distance metrics reflect the beta-diversity of a microbiome sample set based on genomic contents rather than the actual expressed functions.

Here, we developed a novel computational pipeline termed **Phylo**genetically-informed **Func**tional (PhyloFunc) distance to address the above issue by integrating evolutionary relationships with functional attributes to generate functional dissimilarity distances between metaproteomes. We applied PhyloFunc distance to a toy dataset and two real metaproteomic case datasets to evaluate its performance. The results demonstrate that PhyloFunc can group metaproteomes exhibiting functional compensatory behavior between phylogenetically related taxa more closely. Additionally, this approach proved more sensitive to specific environmental responses that were undetectable using other beta-diversity metrics. Finally, we developed a Python package of *PhyloFunc* to implement and streamline the calculation of the PhyloFunc distance algorithm. In addition to supporting custom phylogenetic trees, the package includes an embedded UHGG tree, enabling users to bypass tree input when their protein group IDs are based on the UHGG database [18].

## Results

### The algorithm of PhyloFunc distance

Consider a microbiome sample set as a metacommunity of a total of S species. Metaproteomics analysis can be performed on each of the samples, and a phylogenetic tree of the S species in the metacommunity can be obtained using data from metagenomics, 16S rRNA gene sequencing, or by subsequently retrieving the 16S rRNA gene sequences from databases after inferring taxonomy from metaproteomics data [19]. A phylogeny-informed taxon-function dataset can therefore be summarized (**Fig. 1A**).

**Fig. 1.**
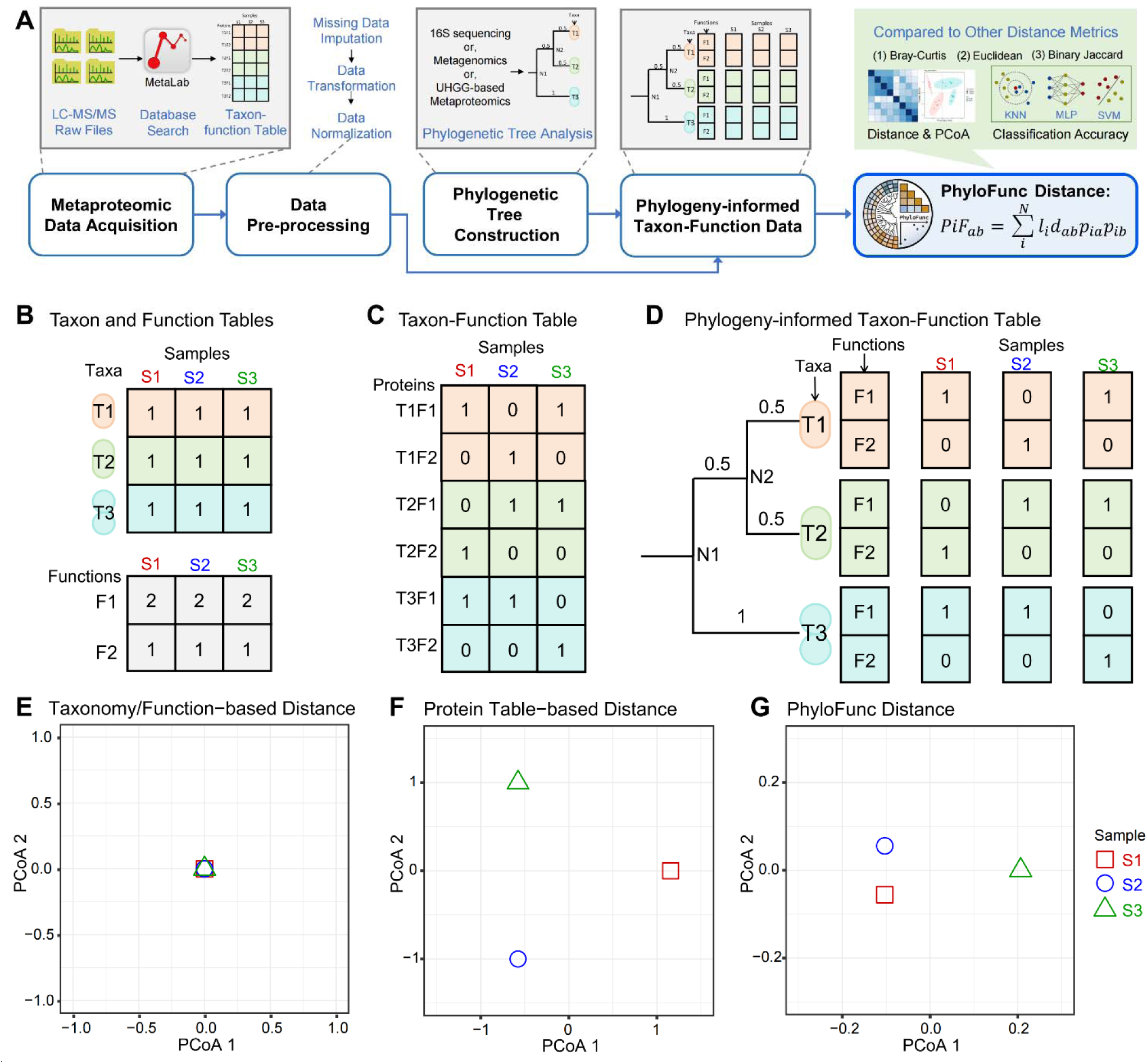
Overview of the PhyloFunc distance metric. **A** LC-MS/MS *.RAW files were acquired, and database search and annotation were performed to obtain taxon-function information of proteins. Through data preprocessing and construction of the phylogenetic tree, phylogeny-informed taxon-function data can be gathered. PhyloFunc distances were computed and compared with other metrics (Bray-Curtis, Jaccard, Euclidean) using Principal Coordinates Analysis (PCoA) [20] and machine learning classification methods (K-Nearest Neighbors Algorithm (KNN) [21], Multilayer Perceptron (MLP) [22], Support Vector Machine (SVM) [23]). **B** Taxon-based or function-based tables of the three microbiome samples. **C** Taxon-function table of the three microbiome samples. **D** Phylogeny-informed taxon-function table. **E** PCoA result of Bray-Curtis dissimilarity based on the taxon-only or function-only table. **F** PCoA result of Bray-Curtis dissimilarity based on taxon-function table. **G** PCoA result of the PhyloFunc distance.

PhyloFunc distance can therefore be computed as the summary of between-sample functional distance of each phylogenetic tree node which is weighted by taxonomic abundance and branch length of the tree. We define PhyloFunc distance *PiF_ab_* between two microbiome samples *a* and *b* as follows:

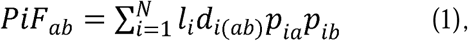

where, *N* is the total number of nodes of the phylogenetic tree (*N* ≥ 5), l is the branch length between node *i* and its ‘parent’, *p_ia_* and *p_ib_* represent the relative taxonomic abundance of samples *a* and *b* at node *I. d_(ab)_* is the metaproteomic functional distance of node *i* between samples *a* and *b* measured by the weighted Jaccard distance between proteomic contents of taxon *i*:

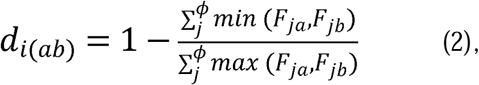

where *ϕ* denotes the total number of functions, *F_ja_* and *F_jb_* represent the normalized functional abundance of the *j* th function in samples *a* and *b*, respectively. A more detailed explanation of the calculation process of the PhyloFunc distance is provided in the **Methods** section, as well as in a step-by-step demonstration in **Supplementary Figures S1-S2**.

We argue that PhyloFunc is a highly informative metric incorporating hierarchical information of taxon-specific functionality and phylogeny of functions. This contrasts with taxon-only, function-only, or taxon-function table metrics, each of which overlooks important relationships between features. First, we demonstrate the strength of PhyloFunc in accounting for the evolutional relatedness of functions in a synthetic toy dataset. This toy dataset is comprised of three samples, each containing six proteins. These proteins are “annotated” to three different taxa and two different functions (**Fig. 1B-D**). The phylogenetic tree specific to the dataset indicates that taxon T1 and T2 are more closely related. First, if we only consider the taxonomic or functional abundances of the metaproteomes, we can sum up protein abundances to obtain taxon-only or function-only tables as in **Fig. 1B**. Another common approach involves calculating protein-level functional distances, where proteins are represented as taxon-specific features (**Fig. 1C**). Finally, we introduce the phylogeny-informed taxon-function dataset as would be required for PhyloFunc, as shown in **Fig. 1D**.

In the toy dataset, by considering only one dimension—either taxonomic or functional abundances—we can design an extreme scenario where the combined profiles of all three samples are identical. Naturally, in such a case, the distances between the samples calculated from taxon-only and function-only datasets are zero (**Fig. 1E**), indicating that taxon-only and function-only data may not capture variability among the samples under certain circumstances. Next, based on the taxon-function dataset, we observed that distances between sample pairs were consistently identical (**Fig. 1F**). In other words, samples are uniformly different from each other when assuming that each protein is equally significant and functions independently. However, as we complemented the dataset with a phylogenetic tree which contains five nodes (including 3 leaves and 2 internal nodes) and simulated weights of branches N2T1 and N2T2 as smaller than N1T3 (i.e., the genetic dissimilarity between T1 and T2 is less than that between T1 or T2 and T3), phylogeny-informed taxon-function data were integrated, enabling the computation of the PhyloFunc distance (details illustrated in **Supplementary Figures S1-S2**). The distance between samples S1 and S2 became closer, while the distance between samples S1 (or S2) and S3 became further apart (**Fig. 1G**). Functional compensation occurs when taxonomically related species undergo functional alterations that allow them to maintain ecosystem processes despite changes in the species’ own functionality. This demonstrates that PhyloFunc sensitively reflects this mechanism as a functional compensation was designed to present between S1 and S2 in the toy dataset.

### Proof-of-principle of PhyloFunc using a synthetic mouse gut microbiome dataset

We next demonstrate that the result presented with our synthetic toy dataset can be true in real-world microbiomes, by analyzing a metaproteomic dataset of mouse gut microbiomes [24]. These mice were inoculated with a synthetic consortium consisting of fourteen or fifteen gut bacterial strains (differentiated by the absence or presence of *B. cellulosilyticus*) and subjected to diets containing different types of dietary fibers. This metaproteomic dataset involved mice allocated to two distinct dietary groups: one fed with HiSF-LoFV (upper tertile of saturated fat content and lower tertile of fruit and vegetable consumption), and the other fed with food supplemented with pea fiber (PEFi). The metaproteomic samples collected on the 19th day of feeding, which exhibited the greatest variation between the samples according to Patnode et al. [24], were chosen for our evaluation of the PhyloFunc method. We performed database search and using MetaLab 2.3 [25] based on author-provided database. The full-length 16S rRNA sequences of the fifteen strains were used to construct a phylogenetic tree (**Supplementary Figure S3**), and functional annotation was performed against the eggNOG 6.0 database [26]. Subsequently, we generated the phylogenetic-tree informed taxon-function dataset of this specific dataset (see **Methods** for details).

We compared the performance of PhyloFunc with three abundance-based distance metrics (Euclidean distance, Bray-Curtis dissimilarity, and Binary Jaccard distance) that use taxon-function data tables as input, i.e. the three methods cannot be informed by phylogenetic information. After computing the three conventional distance metrics and PhyloFunc distance across all samples, the PCoA method was used to visualize and reduce the dimensions of the metrics to show the functional beta-diversity between different samples and groups (**Fig. 2A**). For all metrics, we observed clear separations between two diet groups, i.e., the HiSF-LoFV group (represented by brown points) and the PEFi group (purple points). Samples were also distinguished by fourteen-member communities (circle points) and fifteen-member communities (triangle points). The HiSF-LoFV group showed the contrast between fourteen species and fifteen species communities by all four distance metrics. However, the contrast within the PEFi group was much smaller in the PhyloFunc PCoA result, whereas it appeared more pronounced in PCoA plots of the other three metrics.

**Fig. 2.**
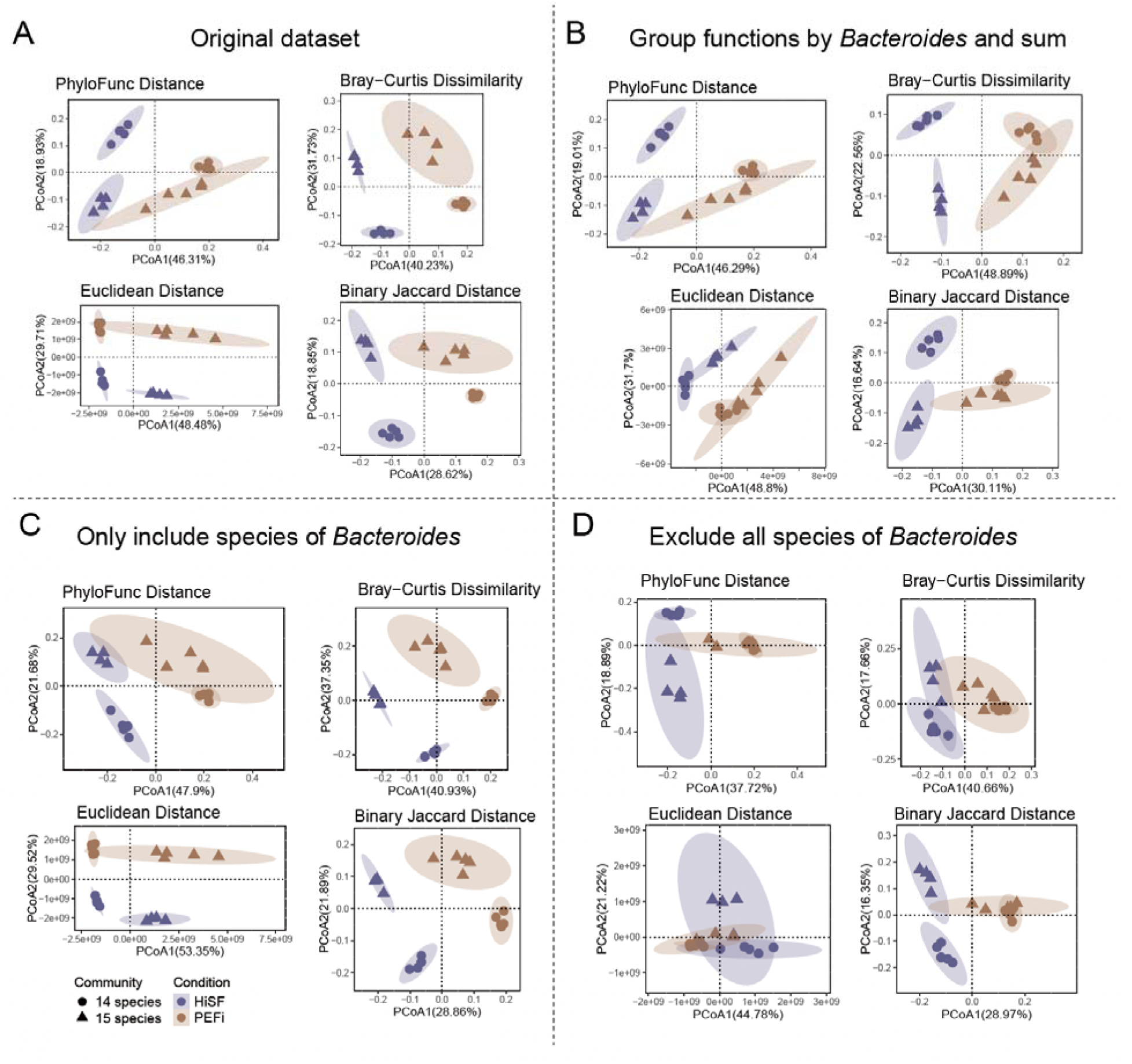
Comparison of different distance metrics using mouse gut microbiome dataset. **A** based on taxon-function table without incorporating phylogenetic information (Euclidean, Bray-Curtis, Jaccard) and based on taxon-function table informed by phylogenetic tree (PhyloFunc). **B** the four distance metrics are based on the sum of functions across *Bacteroides* species. **C** The four distance metrics are based solely on the species of *Bacteroides*. **D** the four distances metrics based on the species excluding *Bacteroides*.

To explore the underlying ecological origination of this phenomenon, we first aggregated each function across the seven *Bacteroides* and *Phocaeicola* species (i.e., *B. caccae*, *B. cellulosilyticus*, *B. finegoldii*, *B*. *ovatus*, *B. thetaiotaomicron*, *P. massiliensis*, *P.vulgatus*) to form a “*Bacteroides*” supergroup, while maintaining the functional profiles of the other eight taxa unchanged. This outcome is reflected in the PCoA plots shown in **Fig. 2B**, we observed that the result corresponding to PhyloFunc was similar to that of the original dataset, whereas PCoA results from the other three distance measures display reduced distances between the two types of communities fed with PEFi. This indicates that PhyloFunc distance effectively recognizes the functional compensatory effect of *Bacteroides*, whereas other distance measures may magnify the impact of functional differences between *Bacteroides* on ecosystem functionality. Finally, to further validate this observation, we calculated these four distances separately for *Bacteroides*-specific data (**Fig. 2C**) and data excluding *Bacteroides* species **(Fig. 2D)** before implementing the PCoA analyses. The results showed that PCoA plots based on *Bacteroides* (**Fig. 2C**) closely resemble those obtained from the original dataset, maintaining a distinct separation between the two PEFi communities across three conventional methods. However, when all *Bacteroides* data were excluded **(Fig. 2D)**, the three conventional PCoA plots exhibited clustering outcomes similar to PhyloFunc distances calculated from grouped *Bacteroides* functions. This indicates that, when features were considered independent in this dataset, *Bacteroides* played a predominant role in shaping the PCoA outcome. In contrast, PhyloFunc demonstrates its capability for hierarchical management of functional alterations among taxonomically related species by weighing functional dissimilarities between these taxa with smaller branch lengths.

Since PhyloFunc is derived from the original UniFrac concept but replaces taxonomic intensit differences with metaproteomic functional distances at the nodes, we further compared it with UniFrac (**Supplementary Figure S4**). The results showed that UniFrac failed to achieve clear separations in PCoA, further demonstrating that PhyloFunc provides superior resolution by integrating functional dimensions. This highlights its advantage in capturing microbial community dynamics more effectively.

### PhyloFunc exihibits sensitivity to *in vitro* human gut microbiome drug responses

To further demonstrate the effectiveness of PhyloFunc distance, we applied our PhyloFunc metric to a more complex multi-dimensional dataset from a live human gut microbiome exposed *in vitro* to different drug treatments [27]. The experiments were performed using the RapidAIM assay [28]. In this experiment, a human gut microbiome sample was subjected to five different drugs - azathioprine (AZ), ciprofloxacin (CP), diclofenac (DC), paracetamol (PR), and nizatidine (NZ). These drugs were administered at three distinct concentrations: low (1000μM), medium (5000μM), and high (biologically relevant drug concentrations as reported by Li et al, 2020 [28]), three technical replicates were performed for each treatment. We reanalyzed the dataset using a database generated by metagenomic sequencing of the microbiome’s baseline sample and performed metagenomic tree construction and taxon-function table preprocessing (see **Methods**). Taxonomic and functional annotations resulted in a taxon-function table containing 973 OGs and 99 genera. The phylogenetic tree constructed by a Maximum Likelihood method comprises 228 nodes (including 115 leaf nodes), along with the calculated weights for 228 branches (**Supplementary Figure S5**).

After calculating the PhyloFunc distance and the other three distances based on preprocessed data, hierarchical clustering and PCoA were both implemented for each drug to compare the functional analysis ability (**Fig. 3A-C, Supplementary Figure S6-S9**). For all samples, hierarchical clustering results based on the four distance metrics effectively reflected the impact of drugs on the diversity of the gut microbiome. PhyloFunc method showed consistency with other metrics and can effectively group samples corresponding to drugs (**Fig. 3A**). This was particularly evident for drugs CP, DC, and NZ, which had marked effects on microbiome functional profiles. This consistency and effectiveness in grouping further proved the method’s validity and robustness. Furthermore, we observed that samples stimulated with high concentration of PR, which did not show clustered responses at the taxon-function level with the other three distance-based methods, were effectively clustered by the PhyloFunc method.

**Fig. 3.**
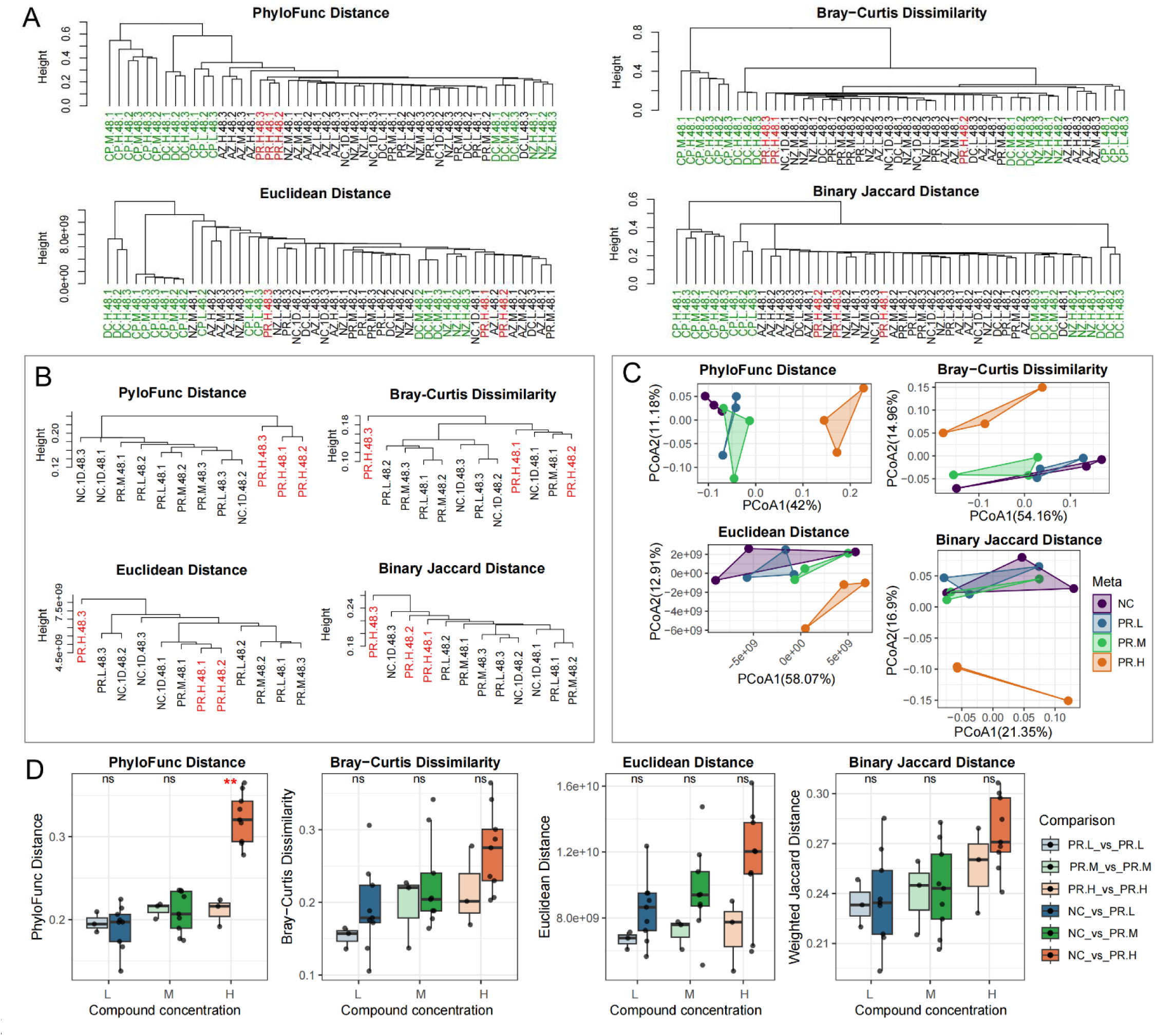
Comparison of different distance metrics on a drug-treated human gut microbiome. **A** Hierarchical clustering of all samples treated with different drugs and concentrations. Green letters indicate that technical triplicates are shown in a cluster, while red letters mark samples treated with high concentrations of paracetamol (PR). **B** Hierarchical clustering methods based on four distances of PR and negative control (NC) samples. **C** PCoA plots based on four distances of drug PR and NC samples. **D** Statistic comparisons based on four distances for drug PR and NC. Each box on the left side of each concentration represents the distribution of technical triplicates within the PR group, indicating the central tendency of distances. Each box on the right side of each concentration depicts the differences between the PR and control groups (NC), with asterisks used to reflect the significance of these differences by two-sided *t*-test.

For a more detailed comparison, we subdivided the data for each drug and compared the effects of each drug group to the control group (NC) on microbial diversity. The results for PR drugs are shown in **Fig. 3B-D**, while those results for other drugs are presented in the Supplementary Materials (**Supplementary Figures S6-S9**). For high concentrations of PR, the PhyloFunc distance method grouped the samples of microbiome showing weak responses to the PR drug compared with the control group (NC). Meanwhile, the PCoA results indicated that the PhyloFunc distance method can distinguish different concentrations of the PR drug from the NC in **Fig. 3C**. However, it was evident from the PCoA results that there were larger overlapping regions between the PR and the NC samples when using the other methods (**Fig. 3C**). We examined the statistical significance of the clustering by comparing distances between replicates to distances between groups (**Fig. 3D**). It can be observed that for high concentration of PR drug, between-group PhyloFunc distance is significantly higher than the between-replicate PhyloFunc distance, which indicates that a drug response has been detected. The other three metrics had no statistical significance in this comparison. For the same set comparisons performed on the other four drugs presented in **Supplementary Figures S6-S9**, it becomes evident that the PhyloFunc method achieves superior or equivalent levels of significance in detecting drug responses.

We further performed PERMANOVA tests using the human gut microbiota datasets, analyzed the effects of different compounds separately and assessed the differential separation of groups across varying concentrations (**Supplementary Tables S3-S7**). Results showed that within the compound groups PR, NZ, DC, and CP, PhyloFunc demonstrated the lowest p-value among all four metrics, or at least equivalent to one other metric in one case. Whereas one exception with AZ shows the lowest p-value with Binary Jaccard distance, while PhyloFunc still showing PERMANOVA significance along with highest R2 and F-values across all four distance measurements.

Despite the overall merits of PhyloFunc over other metrics shown in this dataset, we argue that its strength does not lie in achieving the greatest discrimination among groups compared to other metrics.

Instead, it stands out in integrating phylogenetic and functional information to provide deeper ecological insights, which can sometimes manifest as sample discrimination, as demonstrated in this case.

### PhyloFunc shows higher predictive power in analyzing microbial responses

To further evaluate the predictive power of PhyloFunc in comparison to conventional distance metrics, we employed classification algorithms (KNN, MLP, SVM) to construct machine learning models. These models were built based on four different distance metrics to predict the identity of drugs. Due to the limited sample size, leave-one-out cross-validation was applied to evaluate the accuracy of different models [29, 30] and to compare their performance, resulting in the accuracy comparison results as depicted in **Fig. 4**. For each classification algorithm, we fine-tuned the parameters as detailed in **Supplementary Table 2**. All the three algorithms predicting classification performance showed that PhyloFunc resulted in higher, if not equivalent, predicted accuracy compared to those based on the other three distance metrics.

**Fig. 4.**
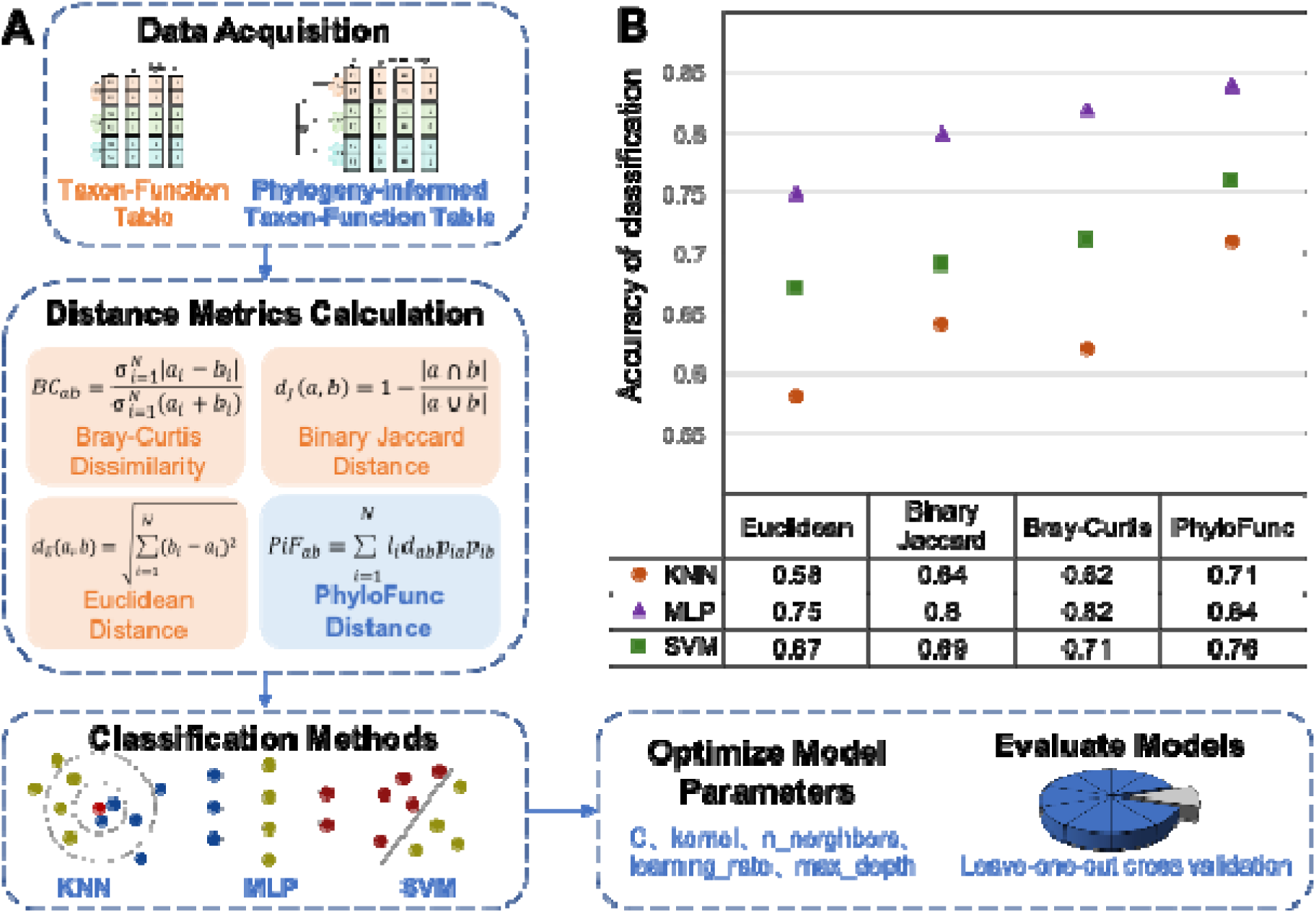
Evaluating accuracies of different distance metrics using three classification algorithms. **A** Data acquisition for a phylogeny-informed taxon-function table, distance metrics calculation, classification methods, and evaluation models. **B** The accuracy classification evaluation of Bray-Curtis dissimilarity, Binary Jaccard distance, Euclidean distance, and PhyloFunc distance metrics based on KNN, MLP, and SVM algorithms.

### Streamlined functional distance calculation with the *PhyloFunc* package

To enhance the broad applicability of PhyloFunc, we have developed a user-friendly Python package of *PhyloFunc* (https://pydigger.com/pypi/PhyloFunc), which includes two primary functions: *PhyloFunc_distance()* for calculating the distance between a pair of samples, and *PhyloFunc_matrix()* for computing a distance matrix across multiple samples. The package offers flexibility with two input options for phylogenetic trees. First, users can provide custom phylogenetic trees in Newick format, constructed from their sample-specific metagenomics or 16S rRNA gene amplicon sequencing data, enabling broader applicability for various research contexts (as we have demonstrated using the synthetic mouse gut microbiome dataset and the in vitro human gut microbiome datasets, **Fig. 2** & **Fig. 3**). Second, **i**t incorporates an embedded phylogenetic tree (bac120_iqtree_v2.0.1.nwk) from the UHGG database[18] as the default input, enabling users to bypass sequencing metagenomic data when their metaproteomic search relies on the UHGG database. We compared the results between UHGG-based and metagenomics-based trees and show highly reproducible results (**Supplementary Figure S10** versus **Figure 3A**). To further support users, we have provided a step-by-step tutorial on GitHub (https://github.com/lumanottawa/PhyloFunc/tree/main/0_PhyloFunk_package_tutorial), which includes detailed instructions, example input file formats, and implementation guidelines. This comprehensive package and its accompanying resources are designed to remove barriers to computational analysis with PhyloFunc, enabling researchers, including those without a bioinformatics background, to easily integrate it into their metaproteomics and microbiome studies.

## Discussion

Metaproteomics is an informative approach to studying the functionality of the human gut microbiome, and its implications in human health and disease. Evaluation of beta-diversity is often one of the initial steps in metaproteomics data exploration. However, there has been a lack of a measurement tool that effectively captures the ecology-centric variations in metaproteomics data. The beta-diversity of gut metaproteome samples is influenced not only by the abundance of taxa and taxon-specific functional compositions but also by the phylogenetic relatedness between taxa. Therefore, including phylogenetic information with protein group taxonomic and functional annotations can better empower researchers to explore both the functional and ecological dynamics of microbial communities, offering insights much overlooked by solely considering taxonomic and functional abundances.

Here, we proposed a novel beta-diversity metric, PhyloFunc, which provides a comprehensive perspective to better detect functional responses to drugs by incorporating phylogenetic information to inform functional distances. Through a simulated dataset, we illustrated the calculation process and indication of the PhyloFunc distance method. This simple toy dataset makes it possible for readers to follow the calculations and understand the hierarchy of PhyloFunc algorithm more effectively. It hierarchically incorporates functional abundance of proteins, taxonomic abundance, and phylogenetic relationship between taxa. As demonstrated by the proof-of-concept toy dataset, as well as a real-world dataset, we report that PhyloFunc distance can account for the functional compensatory effect among taxonomically related species, and offered a more ecologically relevant measurement of functional diversity compared to the three established distance methods tested. Functional compensation can mitigate the impact of species loss or functional changes on the overall ecosystem function, thereby helping maintain ecosystem functions. Research has shown that functional compensation among closely related species with harboring functional redundancy is a key mechanism in sustaining ecosystem functions in response to environmental stimulants[31, 32]. Our PhyloFunc metric is built on such a mechanism, leveraging the functional roles of related taxa to provide a more ecologically relevant measure of ecological beta-diversity.

Furthermore, we tested PhyloFunc using a dataset of *in vitro* drug responses of a human gut microbiome. We first showed that for drugs exhibiting strong effects, PhyloFunc distance showed agreements with other distance metrics. Interestingly, we further observed that for drugs exerting milder effects, the PhyloFunc method can detect new responses and achieve better classification evaluation results than the other tested distance measures, providing deeper insights into drug-microbiome interactions. This result suggests PhyloFunc’s potential for clinical applications. By offering deeper insights into how various drugs affect the functional ecology of the human gut microbiome, PhyloFunc could be useful in developing personalized medicine approaches [33], optimizing drug therapies, and understanding the microbial basis of drug efficacy and side effects. Apart from drug-microbiome interactions, the PhyloFunc metric has significant potential across an even-broader range of applications. These applications extend to any area where evaluation of microbial ecology responses is required, including but not limited to personalized nutrition, prebiotics/probiotics development, disease diagnostics, etc.

## Conclusions

In this work, we introduce a novel metric PhyloFunc and provide its method of computation. The PhyloFunc metric integrates phylogenetic information with taxonomic and functional data to better capture beta-diversity in gut metaproteomes, offering sensitive insights into microbial ecology responses in health and disease applications. To streamline the calculation of PhyloFunc distances, we developed the Python package **PhyloFunc**, which automates the process of calculating functional distances between sample pairs and generates comprehensive distance matrices for multiple samples. This enables efficient assessment of metaproteomic functional beta-diversity across datasets.

## Methods

### Data preparation

#### Metagenomics data processing, taxonomic and phylogenomic analysis

Total genomic DNA from a human stool sample was extracted FastDNA^TM^ SPIN kit with the FastPrep-24TM instrument (MP Biomedicals, Santa Ana, CA). Sequencing libraries were constructed with Illumina TruSeq DNA Sample Prep kit v3 (Illumina, San Diego, CA, USA) according to the manufacturer’s instructions. Paired-end (100-bp) sequencing was performed with the Illumina NovaSeq 6000 at the Génome Québec Innovation Centre of McGill University (Montreal, Canada).

The raw reads were quality-filtered to remove the adapter and low-quality sequences using fastp v0.12.4 (fastp -q 15 -u 40) [34]. The reads were then mapped to the human (hg38; RefSeq: GCF_000001405.39) and phiX reference genomes, and the matches were removed with the Kraken2 v.2.0.9 package. Metagenome assembly of the quality-filtered non-human reads was processed by MEGAHIT v1.2.9 [35] using the --presets meta-large --min-contig-len 1000 parameters. For metagenomic binning, the single_easy_bin command of SemiBin v1.5.1 [36] was used. The resulting bins were then assessed for contamination and completeness with DAS Tool v1.1.4 [37], retaining only high-quality bins or metagenome-assembled genomes (MAGs) with <50% completeness.

The assembled contigs were then annotated using the PROkaryotic Dynamic programming Gene-finding ALgorithm (Prodigal) v2.6.3 [38] to predict open reading frames (ORF). The contigs were translated into amino acid sequences using the anonymous gene prediction mode (prodigal -p meta) and default parameters. The final 115 MAGs were taxonomically classified using the GTDB-Tk v2.1.0 with the r207_v2 [39]. For the phylogenomic analysis, a maximum-likelihood (ML) tree was constructed de novo using the protein sequence alignment produced by GTDB-Tk. First, the aligned sequences were trimmed using trimAl v1.4.rev15 [40] with the heuristic “-automated1” method, and the ML tree was constructed using the IQ-TREE multicore version 2.2.0.3 COVID-edition [41] with 1000 bootstrapping and visualized and annotated using the Interactive Tree of Life (iTOL) web tool [42]. Lastly, the protein-coding sequences of the MAGs were compiled into a single FASTA file and used as the metagenome-inferred protein database for the metaproteomic search.

#### 16S rRNA data processing

The full-length 16S rRNA sequences of the fifteen bacterial strains which consistently colonize animals (**Supplementary Table 1**) were used to construct a phylogenetic tree. Using the Maximum Likelihood method in MEGA v11 [43] with 1000 bootstrapping and default parameters.

#### Metaproteomes database search, taxonomic, and functional annotations

Metaproteomic database searches of the mouse gut microbiome data obtained from Patnode et al., 2019 [24] were performed using MetaLab 2.3 based on the author-provided database of the manuscript (patnodeCommunity_Mmus_allDiets_plus_contams_FR.fasta) with default parameters. Briefly, search parameters included a PSM FDR of 0.01, protein FDR of 0.01, and site FDR of 0.01. Minimum peptide length was set to 7. Modifications considered in protein quantification included N-terminal acetylation and methionine oxidation. The analysis also utilized matching between runs with a time window of 1 minute. For taxonomic annotation, we used the protein names from the fasta file headers of the author-provided database to infer taxonomic originations of the proteins. Metaproteomic database search of the RapidAIM cultured human gut microbiome was performed using MaxQuant 1.6.17.0 using the sample set specific metagenomics database (protein-coding sequences of the MAGs), and the match between runs option was enabled for label-free quantification with default parameters same as the mouse gut microbiome dataset stated above.

Taxonomic annotation of the synthetic mouse gut microbiota and human gut microbiome datasets was performed in two consecutive steps. First, taxonomic information was extracted by matching protein IDs to their origins: for the mouse microbiome dataset, protein IDs were matched to species-specific protein IDs based on their taxonomic origins; for the metagenomic MAG database-searched human gut microbiome dataset, protein IDs were matched to the metagenomic MAG taxonomic classification results. Second, lowest common ancestors (LCAs) were generated at the protein group level, and species-level LCAs were subsequently extracted for further analysis. The datasets achieved 88% and 73% species-level LCA matches at the protein group level for the mouse and human microbiome datasets, respectively. Functional annotation was performed against the eggNOG 6.0 database [26] using DIAMOND v2.1.10 [44] with BLASTp [45], applying a stringent e-value threshold of 10^-5^, under a Linux environment. Root-level orthologous groups (OGs) from the top-1 annotation were used for further analysis, resulting in seed ortholog annotation coverage rate of 99.80% ± 0.03% (Mean ± SD, N = 3 datasets), and OG annotation coverage rate of 99.50% ± 0.11% (Mean ± SD, N = 3 datasets).

Furthermore, an additional metaproteomic database search of the same human microbiome was conducted using the UHGG database with Metalab MAG 1.0.7 [46] and quantitative analysis performed by PANDA v1.2.7[47]. The UHGG database contains a phylogenetic tree that can be directly accessed by the PhyloFunc package. The UHGG database includes a phylogenetic tree that is directly compatible with the PhyloFunc package. Moreover, for microbiomes analyzed using the UHGG database, genome IDs (corresponding to tree nodes) can be directly inferred from genome-specific protein IDs. Functional annotation is also performed using eggNOG 6.0.

#### Data preprocessing

From data preparation, we obtained three different data files for each metaproteomic dataset, i.e. a protein group table with abundance information, a taxonomic table, and a functional table with annotation information. First, we filtered out any protein group with the “REV_” indicator in the protein group table, removed contaminant proteins, and included intensities of microbial protein groups based on label-free quantification (LFQ). Based on the taxonomic and functional annotations described above, we aggregated protein abundances by grouping them according to the same functional OG IDs within the same taxonomic lineage. Subsequently, all of the taxa in the taxon-specific functional table were renamed to align with the names of all leaf nodes in the tree file. Simultaneously, the tree was traversed by a recursive method to assign names to all internal nodes to create a branch table. This table included each branch’s information such as precedent, consequent, the number of child nodes, and branch length. For calculating PhyloFunc distances, branch length values were extracted from the branch table, corresponding to the rows in the “consequent” column whose values matched the taxon names in the taxon-function table. For the case of the sum of functions across *Bacteroides* species in the mouse gut microbiome dataset, we utilized a single “*Bacteroides*” node instead of the subtree encompassing all of Bacteroides species. The branch length value for a *Bacteroides* node was 0.04, which was the length of the branch connecting this subtree. To this end, we gathered the phylogeny-informed taxon-function dataset, which was comprised of two components: the taxon-function table and the branches table (similar as illustrated in **Fig. 1D**).

#### The calculation process of the PhyloFunc distance and other traditional distances

Both the construction of the phylogenetic tree and the computation of the PhyloFunc distance were implemented through programming in Python. To illustrate the calculation process of PhyloFunc-based distance most clearly, we employed a simulated dataset as a demonstration (**Supplementary Figures S1-S2)**. Briefly, based on taxon-function data obtained by preprocessing methods, relative abundance of each function within each taxon and the relative abundance of taxon were calculated by taxon-specific protein biomass contributions. Secondly, the relative functional abundance was weighed by their corresponding relative taxonomic abundance and then expanded to represent all nodes up to the root of the phylogeny by summing up each node to get the expanded table. Similarly, the taxonomic table was converted into the expanded table by summing up all nodes in the phylogeny tree. Thirdly, functional distances between each sample pair were calculated according to Eq. (2). Finally, each functional distance was weighed by branch length and relative protein abundances in samples pairs and PhyloFunc distances between samples were then calculated according to Eq. (1). Other methods of Bray-Curtis dissimilarity, Binary Jaccard distance and Euclidean distance were calculated using the R package “ vegan” [35]. For Binary Jaccard distance, we considered non-zero numbers as 1 in binary and used the parameter “binary” to calculate distances between sample pairs.

#### Evaluation and visualization

Different methods including PCoA, statistical tests, hierarchical clustering, classification algorithms and PERMANOVA tests were applied to evaluate the performance across different distances. The detail of the evaluation can be found in the figure legends and main texts. The PCoA analysis was realized by R function dudi.pco() in package ade4. PCoA plots were visualized using the R package ggplot2, with the aspect ratio standardized to 1:1 to ensure a consistent comparison. In the PCoA plots of the human gut microbiome dataset, three replicated points for each group were connected with straight lines and displayed as triangles. Box plots also were visualized using the R package ggplot2. PERMANOVA was performed using R function adonis2(). Hierarchical clustering was performed using the R function hclust() and hierarchical clustering plots were visualized using the R package stats. Based on normalized PhyloFunc and the other three distance metrics, we selected three standard algorithms—KNN, MLP, and SVM—to construct classification models and employed a Leave-One-Out Cross-Validation (LOOCV) approach for splitting training and test sets. The distance matrix was used as the input sample data, with the names of five drugs as classification labels. In each iteration, one sample from the distance dataset was designated as the test set, while the remaining samples were used as the training set to build the classification model. The performance of each model was evaluated by comparing its prediction for the test sample with the true drug classification. Accuracy was calculated as the proportion of correctly classified samples across all iterations. We used the grid search method to obtain the optimal parameters for each classification algorithm of different distance methods. The primary optimal parameters were presented in **Supplementary Table 2**, while the corresponding high-accuracy evaluation results were illustrated in **Fig. 4B**. The classification models were implemented in Python 3.11, and Python packages Pandas, Numpy, sklearn were used. The grid search method for the selection of optimal parameters was implemented in the Python package sklearn.model_selection.

## List of abbreviations

Phylo-RPCA: Phylogenetic Robust Principal-Component Analysis
POMS: Phylogenetic Organization of Metagenomic Signals
UHGG: The Unified Human Gastrointestinal Genome collection
PCoA: Principal Coordinates Analysis
KNN: K-Nearest Neighbors Algorithm
MLP: Multilayer Perceptron
SVM: Support Vector Machine
MAGs: Metagenome-assembled Genomes
ORF: Open Reading Frames
ML: Maximum-likelihood
iTOL: Interactive Tree of Life
LFQ: Label-free Quantification

## Declarations

### Ethics approval and consent to participate

No new participant was recruited in this study as data used was obtained from previous work [20, 23]. Ethics approval for human stool sample collection in the previous work [27] was performed by the Ottawa Health Science Network Research Ethics Board at the Ottawa Hospital (approval number: 20160585-01H) and written consent to participate was signed by the participant.

### Consent for publication

Not applicable.

### Availability of data and materials

The metagenomics FASTQ file in this study is deposited at NCBI BioProject SRR29021656. The metaproteomics data of the mouse gut microbiome was obtained from MassIVE data repository of MassIVE: MSV000082287. The metaproteomics data of RapidAIM cultured microbiome was obtained from a previous study which has been deposited to the ProteomeXchange Consortium [48] via the PRIDE partner repository [49] under accession number <u>PXD024845</u>. Additional data from the analyses presented in this paper are available in the Supplementary Material, and the corresponding visualization and analysis input data and code have been deposited in the GitHub repository (https://github.com/lumanottawa/PhyloFunc).

### Author’s contributions

LW, CMAS, DF and LL conceptualized the study; LW, CMAS, JMS, DF and LL developed the methodology; LW performed formal analysis; LL, CMAS, ZN and JMS provided resources; LW, CMAS, JMS, LL, BS and JY performed data curation; Y.L. and L.W. developed the Python package and tutorial; LW, LL, JMS and DF wrote the original draft; CMAS, ZN, BS and JY reviewed and edited the manuscript draft; LW and LL visualized the data; DF and LL supervised the study. The authors declare that they follow principles of inclusion & ethics in global research. All authors read, revised, and approved the final manuscript.

## Acknowledgements

We thank all members of the Figeys and the Li labs who have contributed ideas.

