## Supplementary Figure S1 for "PhyloFunc: Phylogeny-informed Functional Distance as a New Ecological Metric for Metaproteomic Data Analysis"

**Table of contents**

**Supplementary Figures:**

**Figure S1.** The calculation process of PhyloFunc distance, part 1.

**Figure S2.** The calculation process of PhyloFunc distance, part 2.

**Figure S3.** The phylogenetic tree of the mouse gut microbiome case dataset.

**Figure S4.** Comparison of four UniFrac distance metrics applied to the mouse gut microbiome dataset.

**Figure S5.** The phylogenetic tree of the human gut microbiome case dataset.

**Figure S6.** Comparison different distances for human gut microbiome by PCoA and statistical analysis between Azathioprine (AZ) and control group (NC).

**Figure S7.** Comparison different distances for human gut microbiome by PCoA and statistical analysis between Ciprofloxacin (CP) and control group (NC).

**Figure S8.** Comparison different distances for human gut microbiome by PCoA and statistical analysis between Diclofenac (DC) and control group (NC).

**Figure S9.** Comparison different distances for human gut microbiome by PCoA and statistical analysis between Nizatidine (NZ) and control group (NC).

**Figure S10.** Comparison of different distance metrics on same human gut microbiome dataset searched against the UHGG database.

**Supplementary Tables:**

**Table S1.** List of microbial species used to construct the phylogenetic tree.

**Table S2.** List of primary optimal parameters for classification.

**Table S3.** PERMANOVA results between Paracetamol (PR) and control group (NC).

**Table S4.** PERMANOVA results between Nizatidine (NZ) and control group (NC).

**Table S5.** PERMANOVA results between Diclofenac (DC) and control group (NC).

**Table S6.** PERMANOVA results between Ciprofloxacin (CP) and control group (NC).

**Table S7.** PERMANOVA results between Azathioprine (AZ) and control group (NC).

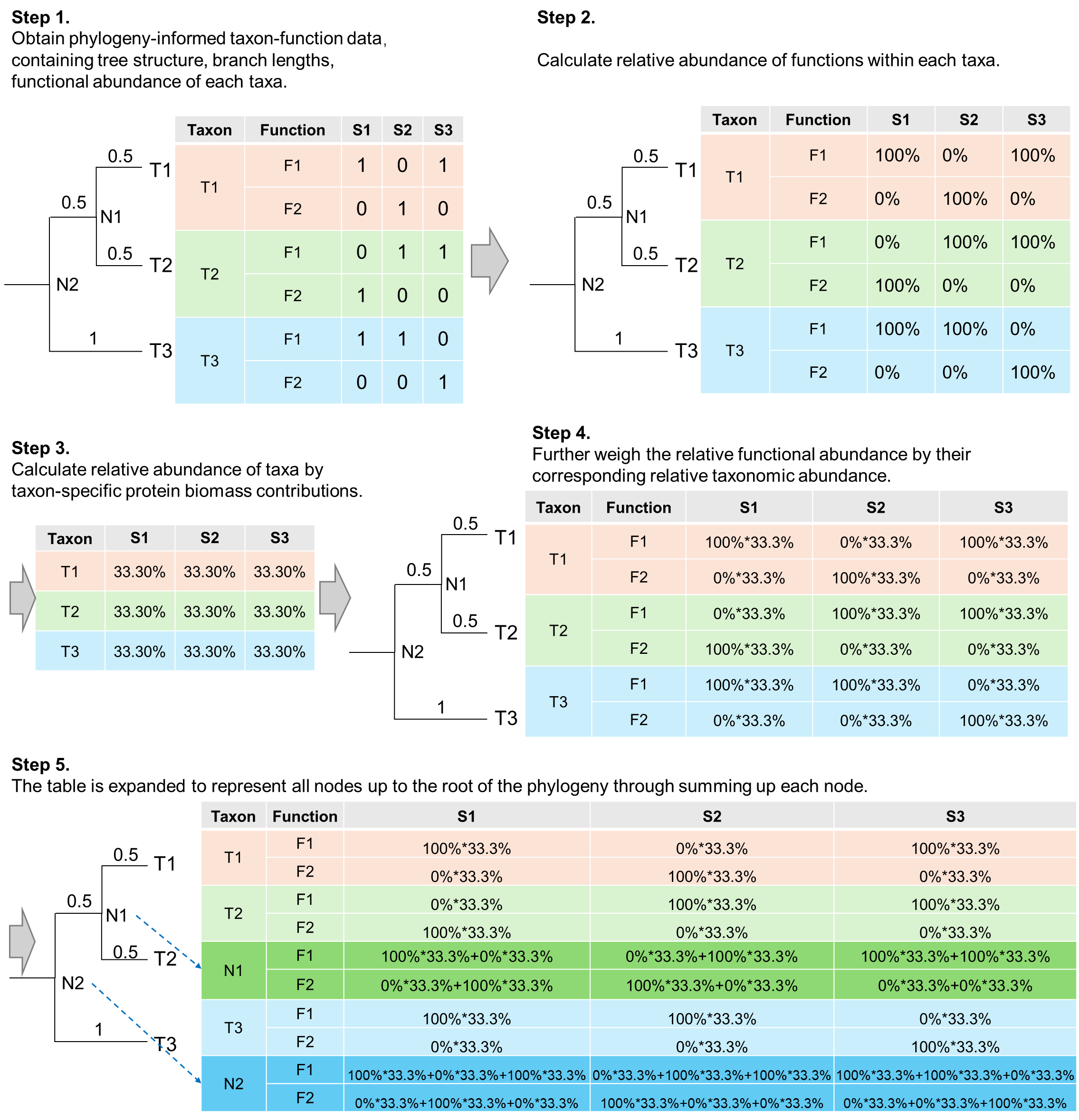

**Figure S1. The calculation process of PhyloFunc distance, part 1.**

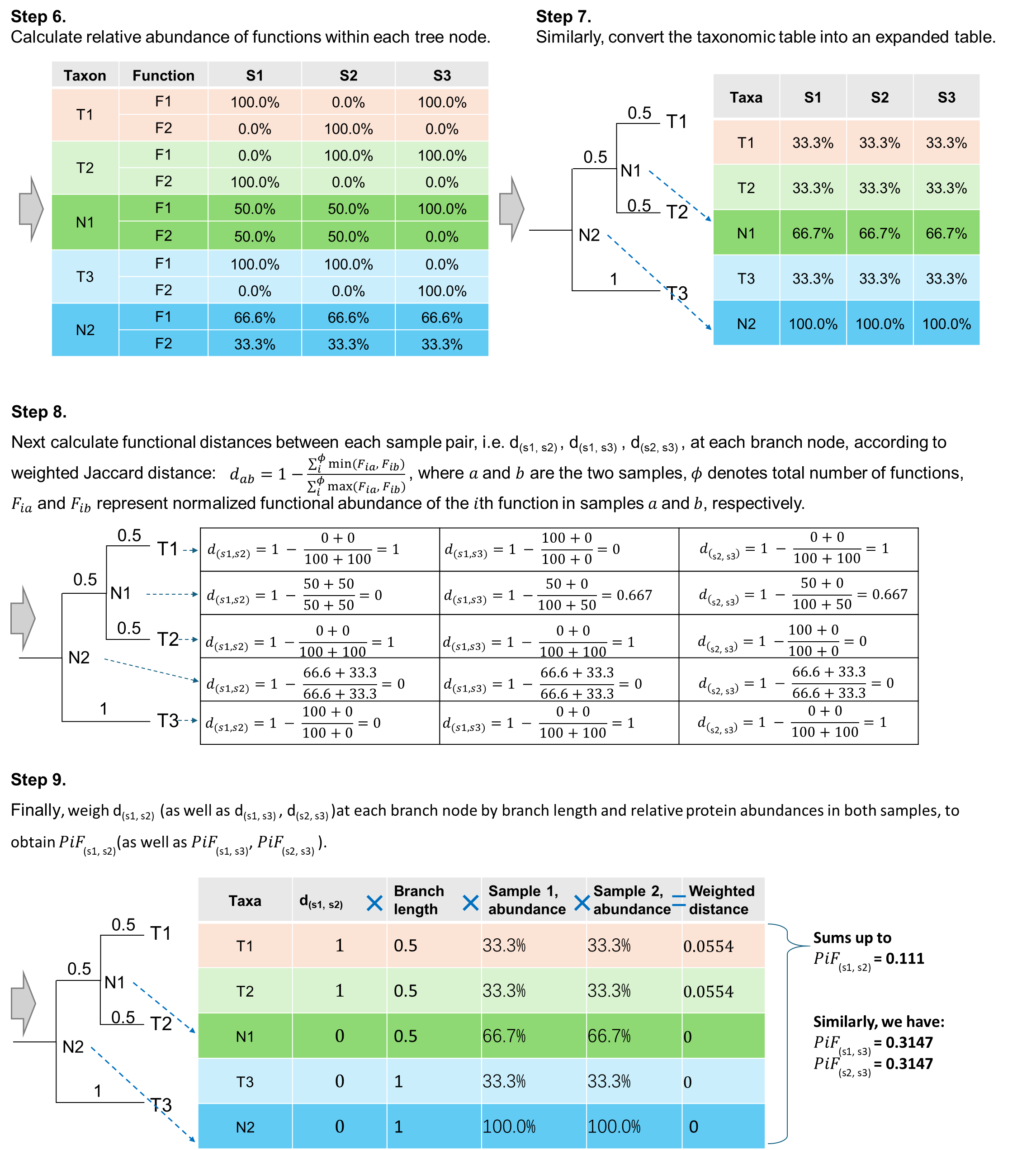

**Figure S2.The calculation process of PhyloFunc distance, part 2.**

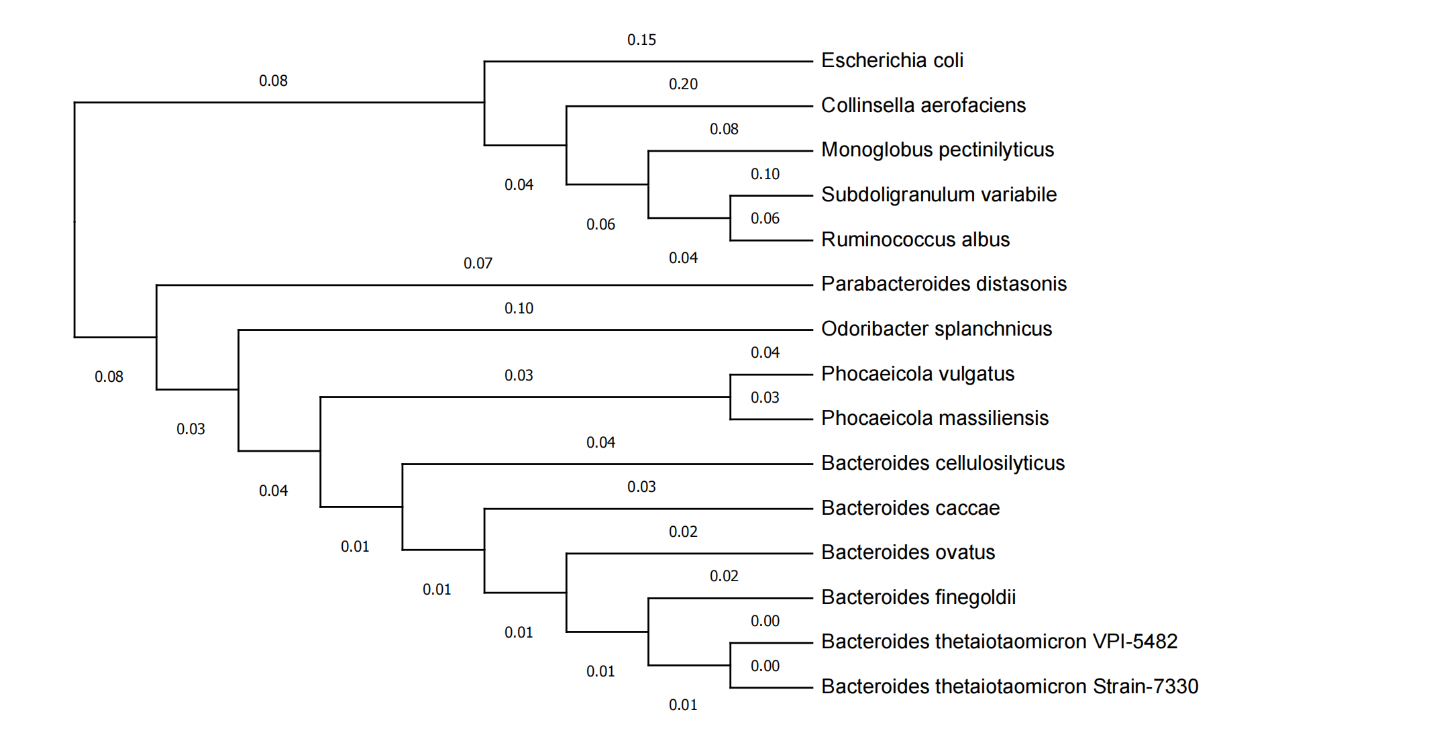

**Figure S3. The phylogenetic tree of the mouse gut microbiome case dataset.**

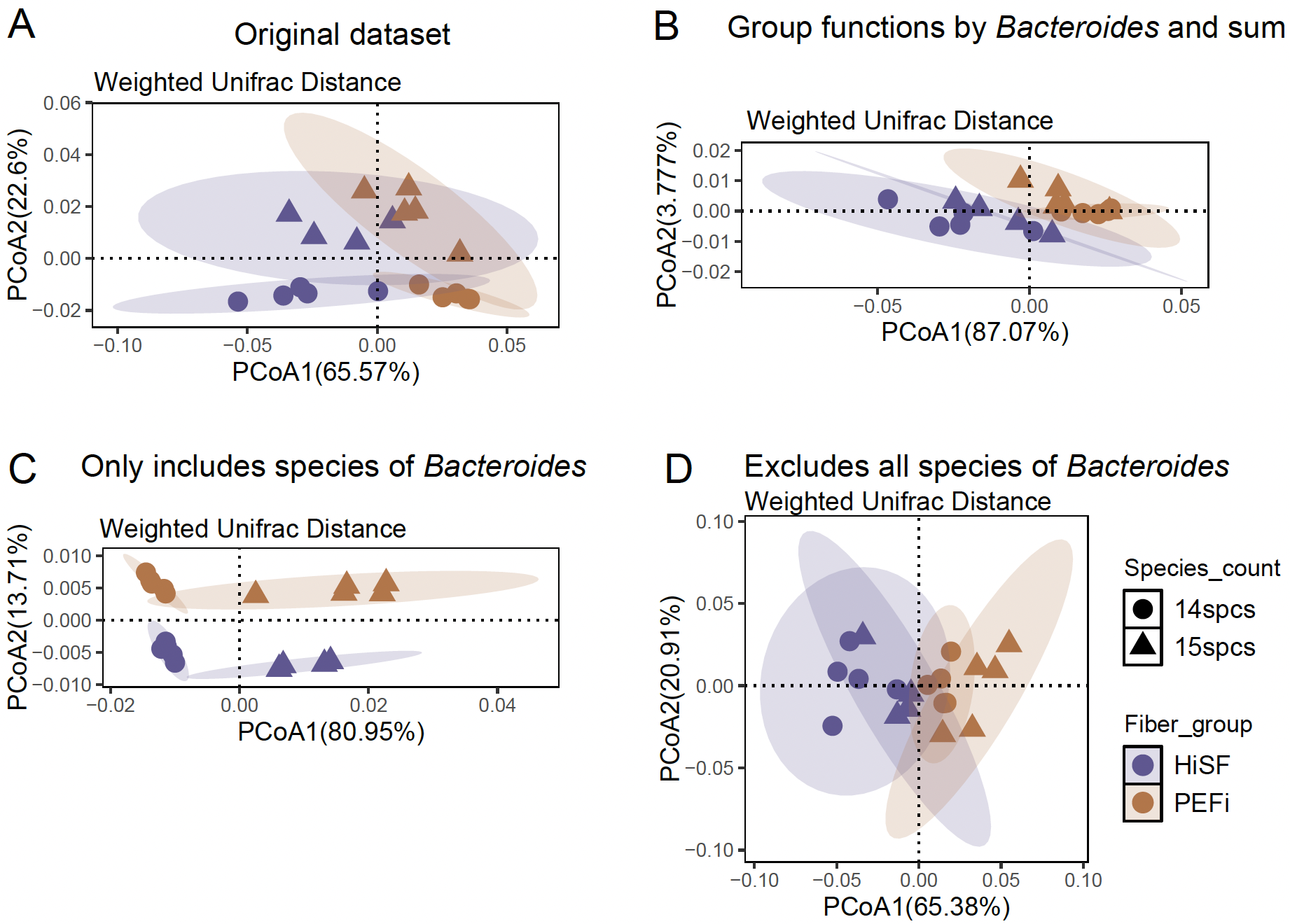

**Figure S4.** **Comparison of four UniFrac distance metrics applied to the mouse gut microbiome dataset.** **A** calculated from the complete taxon intensity table; **B** derived from the functional sum of Bacteroides species; **C** based solely on the species within Bacteroides; and **D**, excluding all Bacteroides species.

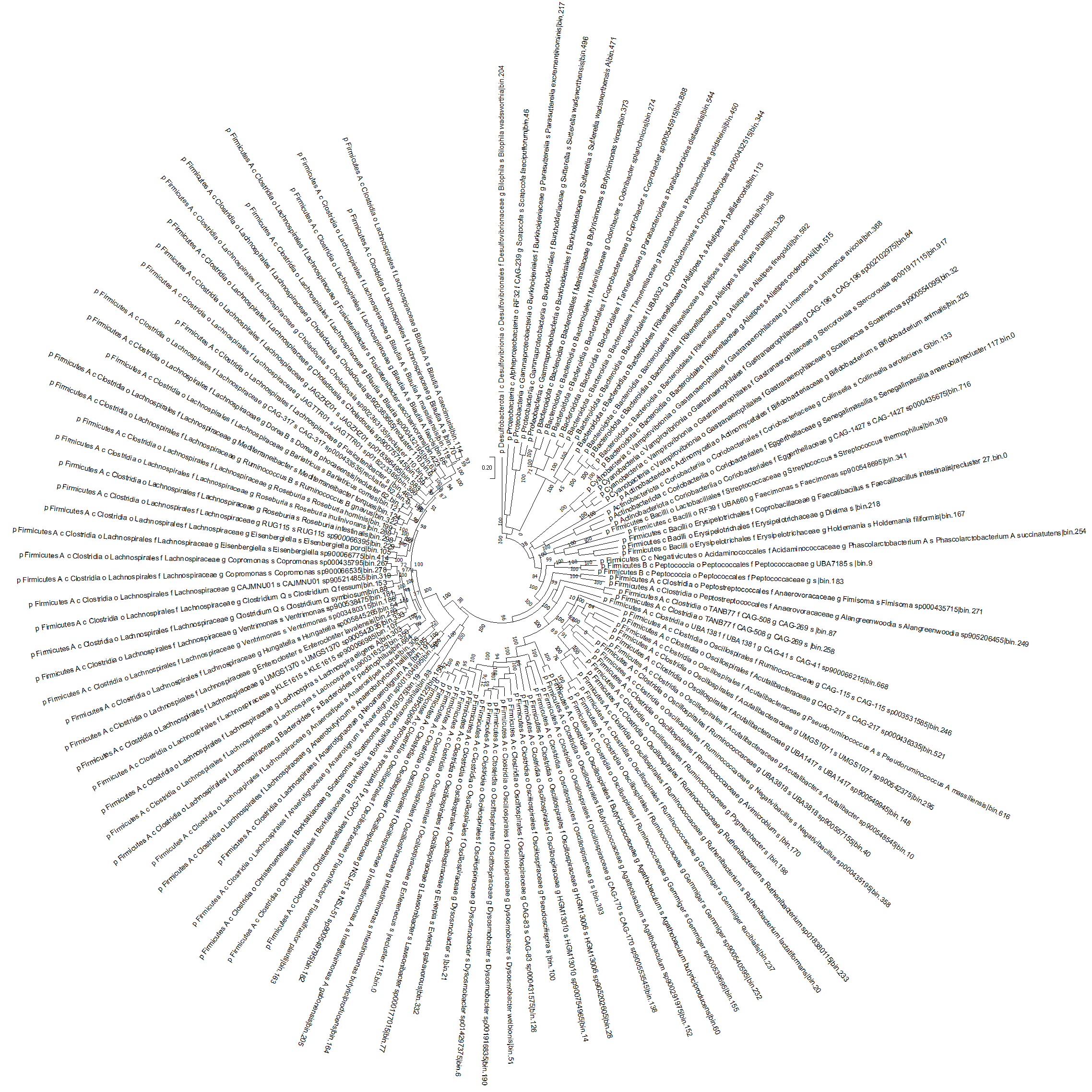

**Figure S5. The phylogenetic tree of the human gut microbiome case dataset.**

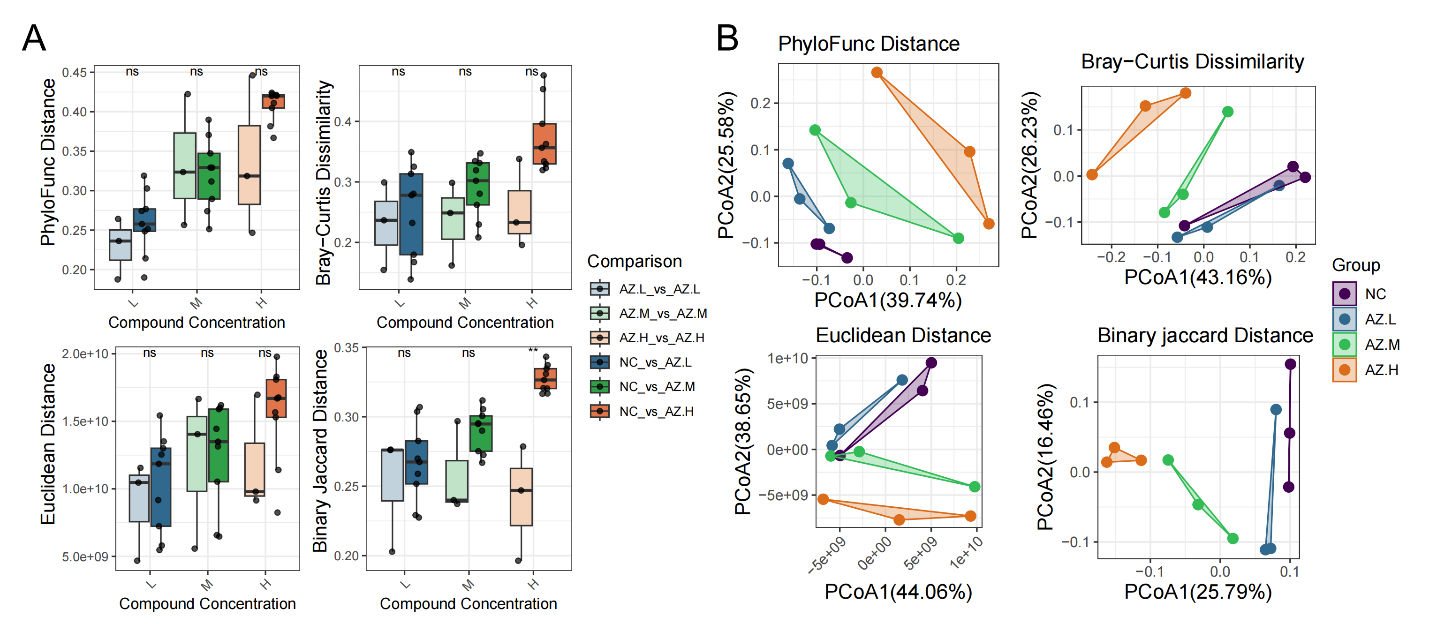

**Figure S6. Comparison different distances for human gut microbiome by PCoA and statistical analysis between Azathioprine (AZ) and control group (NC).**

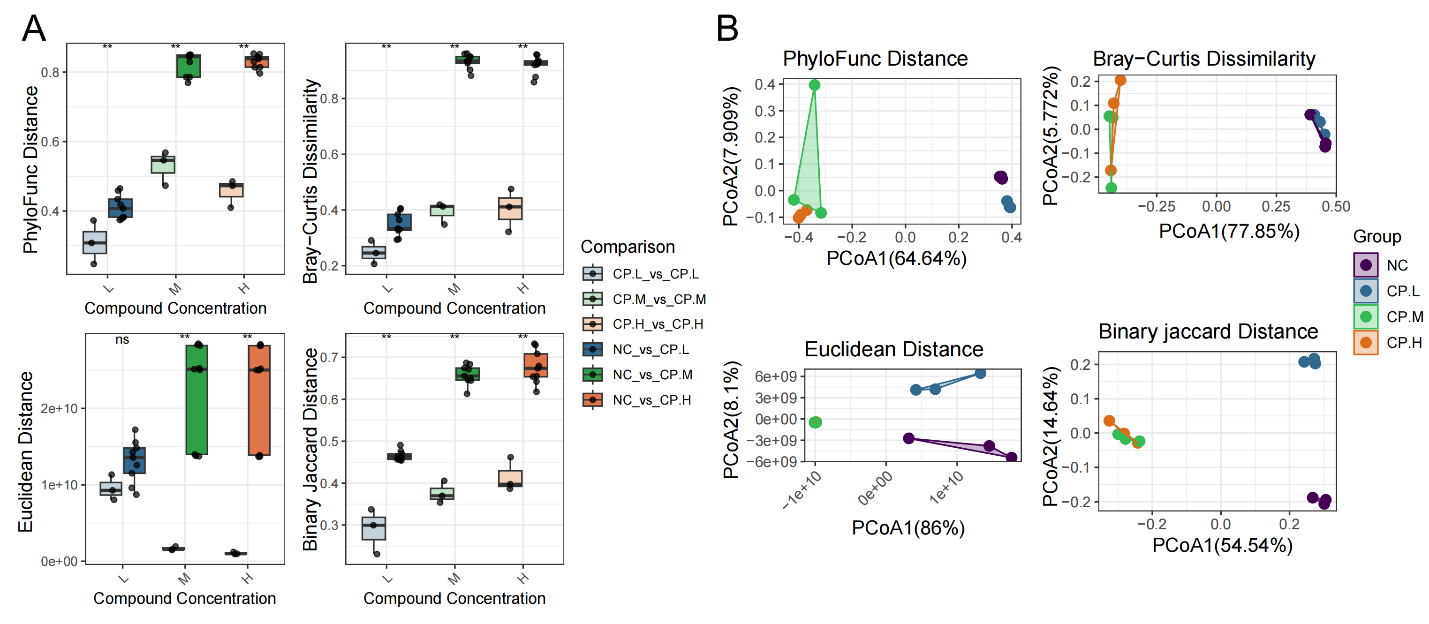

**Figure S7. Comparison different distances for human gut microbiome by PCoA and statistical analysis between Ciprofloxacin (CP) and control group (NC).**

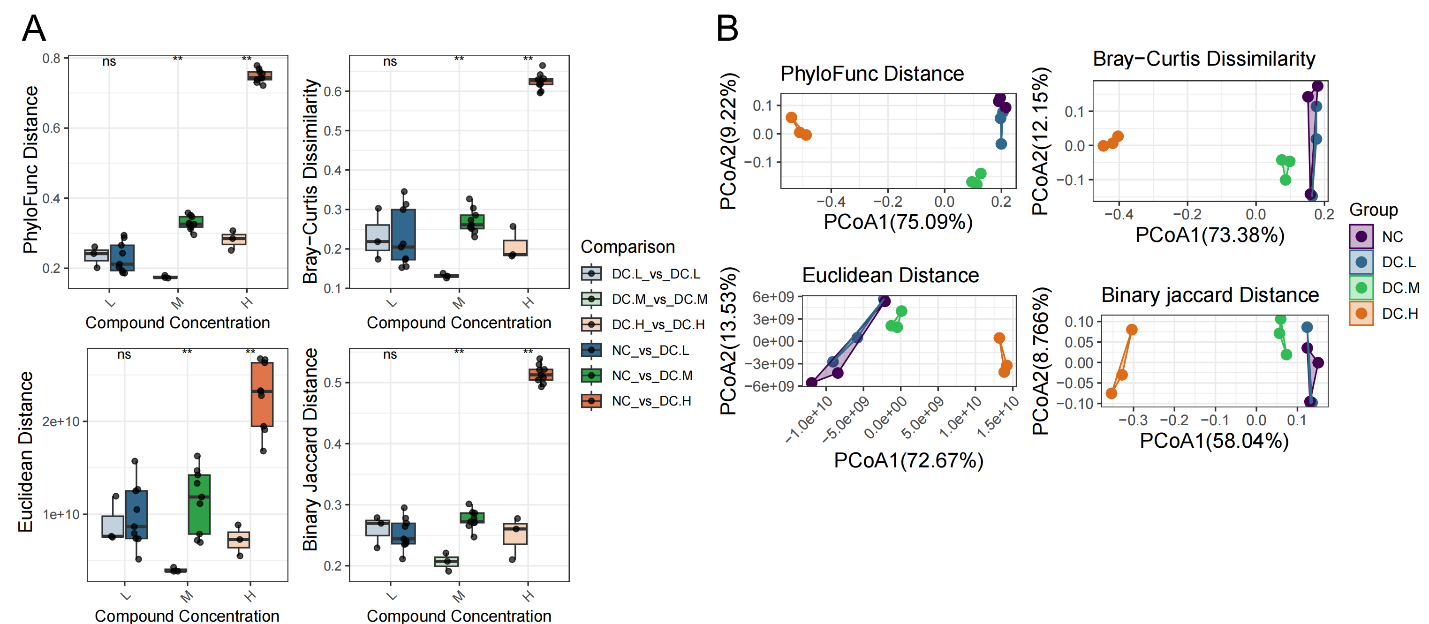

**Figure S8. Comparison different distances for human gut microbiome by PCoA and statistical analysis between Diclofenac (DC) and control group (NC).**

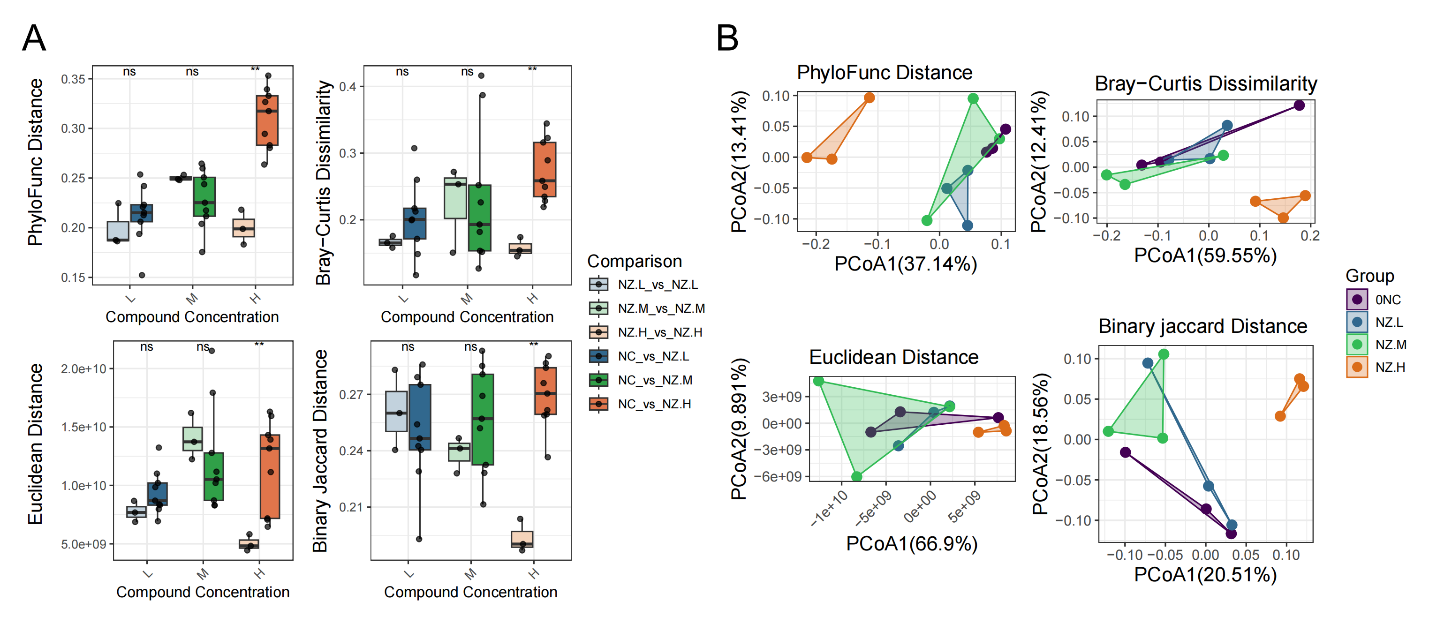

**Figure S9. Comparison different distances for human gut microbiome by PCoA and statistical analysis between Nizatidine (NZ) and control group (NC).**

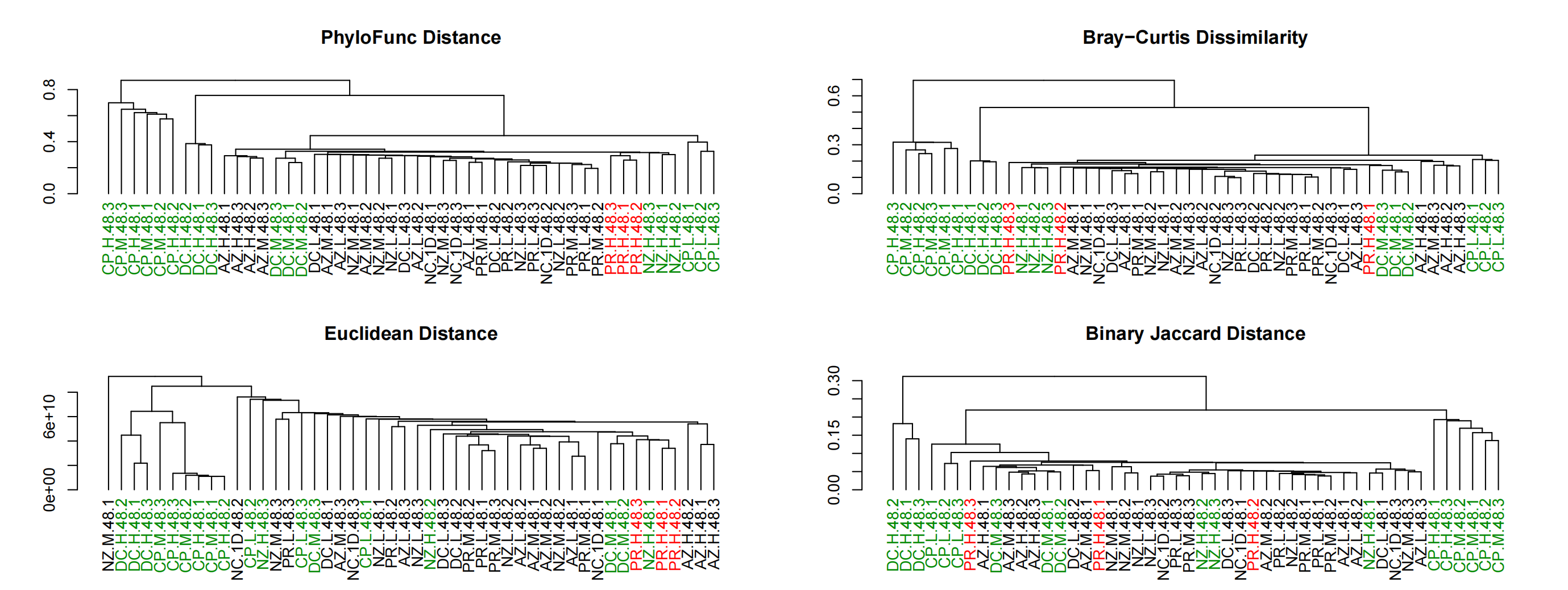

**Figure S10. Comparison of different distance metrics on same human gut microbiome dataset searched against the UHGG database.** Hierarchical clustering of all samples with taxa based on the MGYG (UHGG) database. Green letters indicate that technical triplicates are shown in a cluster, while red letters mark samples treated with high concentrations of paracetamol (PR).

**Table S1. List of microbial species used to construct the phylogenetic tree.**

| **Microbial Community** | **Patnode et al. (2019) [****19****]** | **GenBank Accession** | **16S rRNA bp** | **Link to 16S rRNA gene sequence used for tree** |
| --- | --- | --- | --- | --- |
| *Bacteroides caccae* | *Bacteroides caccae TSDC17.2-1.2* | NR_026242.1 | 1,422 | https://www.ncbi.nlm.nih.gov/nuccore/NR_026242.1?report=fasta |
| *Bacteroides cellulosilyticus* | *Bacteroides cellulosilyticus WH2* | NR_042203.1 | 1,426 | https://www.ncbi.nlm.nih.gov/nuccore/NR_042203.1 |
| *Bacteroides finegoldii* | *Bacteroides finegoldii TSDC17.2-1.1* | NR_041313.1 | 1,485 | https://www.ncbi.nlm.nih.gov/nuccore/NR_041313.1 |
| *Bacteroides ovatus ATCC-8483* |  | NR_040865.1 | 1,455 | https://www.ncbi.nlm.nih.gov/nuccore/NR_040865.1 |
| *Bacteroides thetaiotaomicron 7330* |  | AY895199.1 | 1,345 | https://www.ncbi.nlm.nih.gov/nuccore/AY895199.1 |
| *Bacteroides thetaiotaomicron VPI-5482* |  | NR_074277.1 | 1,483 | https://www.ncbi.nlm.nih.gov/nuccore/NR_074277.1 |
| *Collinsella aerofaciens* | *Collinsella aerofaciens TSDC17.2-1.1/Collinsella aerofaciens TSDC17.2-2.1* | NR_028604.1 | 1,463 | https://www.ncbi.nlm.nih.gov/nuccore/NR_028604.1 |
| *Escherichia coli* | *Escherichia coli TSDC17.2-1.2* | NR_024570.1 | 1,450 | https://www.ncbi.nlm.nih.gov/nuccore/NR_024570.1 |
| *Odoribacter splanchnicus* | *Odoribacter splanchnicus TSDC17.2-1.2* | NR_044636.1 | 1,467 | https://www.ncbi.nlm.nih.gov/nuccore/NR_044636.1 |
| *Parabacteroides distasonis* | *Parabacteroides distasonis TSDC17.2-1.1* | NR_041342.1 | 1,488 | https://www.ncbi.nlm.nih.gov/nuccore/NR_041342.1 |
| *Phocaeicola massiliensis* | *Bacteroides massiliensis TSDC17.2-1.1* | NR_042745.1 | 1,493 | https://www.ncbi.nlm.nih.gov/nuccore/NR_042745.1?report=fasta |
| *Phocaeicola vulgatus ATCC-8483* | *Bacteroides vulgatus ATCC-8483* | NR_074515.1 | 1,510 | https://www.ncbi.nlm.nih.gov/nuccore/NR_074515.1 |
| *Monoglobus pectinilyticus* | *Ruminococcaceae TSDC17.2-1.2* | NR_159227.1 | 1,493 | https://www.ncbi.nlm.nih.gov/nuccore/NR_159227.1 |
| *Ruminococcus albus* | *Ruminococcus albus TSDC17.2-1.1/Ruminococcus albus TSDC17.2-1.4* | NR_025929.1 | 1,407 | https://www.ncbi.nlm.nih.gov/nuccore/NR_025929.1 |
| *Subdoligranulum variabile* | *Subdoligranulum variabile TSDC17.2-1.1* | NR_028997.1 | 1,428 | https://www.ncbi.nlm.nih.gov/nuccore/NR_028997.1 |

**Table S2. List of primary optimal parameters for classification.**

| **Classification Algorithm** | **The range of parameter values** | **Optimal primary parameters** | **Distance Methods** |
| --- | --- | --- | --- |
| KNN | n_neighbors: from 2 to 10  'metric': 'precomputed',  'algorithm': 'brute' | k=2 | Euclidean Distance |
|  |  | k=2 | Bray-Curtis Dissimilarity |
|  |  | k=3 | Binary Jaccard Distance |
|  |  | k=3 | PhyloFunc Distance |
| MLP | 'hidden_layer_sizes': [(5,), (10,), (15,), (10, 5), (15, 10)],  'activation': ['relu', 'tanh'],  'solver': ['adam', 'sgd'],  'alpha': [0.001, 0.01],  'learning_rate_init': [0.001, 0.01, 0.1] | 'activation': 'tanh',  'alpha': 0.001,  'hidden_layer_sizes': (10, 5), 'learning_rate_init': 0.1,  'solver': 'adam' | Euclidean Distance |
|  |  | 'activation': 'relu',  'alpha': 0.001,  'hidden_layer_sizes': (15,), 'learning_rate_init': 0.01,  'solver': 'adam' | Bray-Curtis Dissimilarity |
|  |  | 'activation': 'tanh',  'alpha': 0.001,  'hidden_layer_sizes': (10,), 'learning_rate_init': 0.1,  'solver': 'sgd' | Binary Jaccard Distance |
|  |  | 'activation': 'tanh',  'alpha': 0.001,  'hidden_layer_sizes': (10, 5), 'learning_rate_init': 0.01,  'solver': 'adam' | PhyloFunc Distance |
| SVM | 'C': [0.1,1,4,6,5,8,7,9,10],  'kernel': ['precomputed'],  'gamma':[0.01,0.001,0.02] | 'C': 9, 'gamma': 0.01 | Euclidean Distance |
|  |  | 'C': 5, 'gamma': 0.01 | Bray-Curtis Dissimilarity |
|  |  | 'C': 4, 'gamma': 0.01 | Binary Jaccard Distance |
|  |  | 'C': 8, 'gamma': 0.01 | PhyloFunc Distance |

| PhyloFunc distance | |  |  |  |
| --- | --- | --- | --- | --- |
| Df | SumOfSqs | R^2^ | F | Pr(>F) |
| 3 | 0.16871034 | **0.49931449** | **2.6593646** | **0.0135** |
| 8 | 0.16917358 | 0.50068551 |  |  |
| 11 | 0.33788392 | 1 |  |  |
| Bray-Curtis dissimilarity | |  |  |  |
| Df | SumOfSqs | R^2^ | F | Pr(>F) |
| 3 | 0.08789169 | 0.31254796 | 1.21239181 | 0.3032 |
| 8 | 0.19331856 | 0.68745204 |  |  |
| 11 | 0.28121025 | 1 |  |  |
| Euclidean distance | |  |  |  |
| Df | SumOfSqs | R^2^ | F | Pr(>F) |
| 3 | 1.5642E+20 | 0.3340671 | 1.33774078 | 0.2596 |
| 8 | 3.118E+20 | 0.6659329 |  |  |
| 11 | 4.6822E+20 | 1 |  |  |
| Jaccard distance | |  |  |  |
| Df | SumOfSqs | R^2^ | F | Pr(>F) |
| 3 | 0.10437987 | 0.29888602 | 1.13680428 | 0.2034 |
| 8 | 0.24484982 | 0.70111398 |  |  |
| 11 | 0.34922969 | 1 |  |  |

**Table S3. PERMANOVA results between Paracetamol (PR) and control group (NC).**

| PhyloFunc distance | |  |  |  |
| --- | --- | --- | --- | --- |
| Df | SumOfSqs | R^2^ | F | Pr(>F) |
| 3 | 0.16958991 | **0.47967117** | **2.45829763** | **0.0059** |
| 8 | 0.1839646 | 0.52032883 |  |  |
| 11 | 0.35355451 | 1 |  |  |
| Bray-Curtis dissimilarity | |  |  |  |
| Df | SumOfSqs | R^2^ | F | Pr(>F) |
| 3 | 0.14715114 | 0.43912944 | 2.08784687 | 0.0787 |
| 8 | 0.18794628 | 0.56087056 |  |  |
| 11 | 0.33509742 | 1 |  |  |
| Euclidean distance | |  |  |  |
| Df | SumOfSqs | R^2^ | F | Pr(>F) |
| 3 | 3.5321E+20 | 0.44129325 | 2.10626061 | 0.095 |
| 8 | 4.4718E+20 | 0.55870675 |  |  |
| 11 | 8.0039E+20 | 1 |  |  |
| Jaccard distance | |  |  |  |
| Df | SumOfSqs | R^2^ | F | Pr(>F) |
| 3 | 0.13040253 | 0.36250231 | 1.51635501 | 0.0131 |
| 8 | 0.2293263 | 0.63749769 |  |  |
| 11 | 0.35972884 | 1 |  |  |

**Table S4. PERMANOVA results between Nizatidine (NZ) and control group (NC).**

| PhyloFunc distance | |  |  |  |
| --- | --- | --- | --- | --- |
| Df | SumOfSqs | R^2^ | F | Pr(>F) |
| 3 | 1.2199808 | **0.85489838** | **15.7112577** | **0.0001** |
| 8 | 0.20706694 | 0.14510162 |  |  |
| 11 | 1.42704775 | 1 |  |  |
| Bray-Curtis dissimilarity | |  |  |  |
| Df | SumOfSqs | R^2^ | F | Pr(>F) |
| 3 | 0.79453707 | 0.79883467 | 10.589428 | 0.0033 |
| 8 | 0.2000831 | 0.20116533 |  |  |
| 11 | 0.99462017 | 1 |  |  |
| Euclidean distance | |  |  |  |
| Df | SumOfSqs | R^2^ | F | Pr(>F) |
| 3 | 9.341E+20 | 0.7466147 | 7.85749048 | 0.0026 |
| 8 | 3.1701E+20 | 0.2533853 |  |  |
| 11 | 1.2511E+21 | 1 |  |  |
| Jaccard distance | |  |  |  |
| Df | SumOfSqs | R^2^ | F | Pr(>F) |
| 3 | 0.52397317 | 0.68628831 | 5.83370722 | 0.0006 |
| 8 | 0.23951524 | 0.31371169 |  |  |
| 11 | 0.7634884 | 1 |  |  |

**Table S5. PERMANOVA results between Diclofenac (DC) and control group (NC).**

| PhyloFunc distance | |  |  |  |
| --- | --- | --- | --- | --- |
| Df | SumOfSqs | R^2^ | F | Pr(>F) |
| 3 | 1.97729603 | 0.75838923 | 8.37036899 | **0.0004** |
| 8 | 0.62993512 | 0.24161077 |  |  |
| 11 | 2.60723116 | 1 |  |  |
| Bray-Curtis dissimilarity | |  |  |  |
| Df | SumOfSqs | R^2^ | F | Pr(>F) |
| 3 | 2.42112291 | **0.83870249** | **13.86593** | 0.0007 |
| 8 | 0.4656253 | 0.16129751 |  |  |
| 11 | 2.88674821 | 1 |  |  |
| Euclidean distance | |  |  |  |
| Df | SumOfSqs | R^2^ | F | Pr(>F) |
| 3 | 1.3363E+21 | 0.83833105 | 13.8279457 | 0.0012 |
| 8 | 2.577E+20 | 0.16166895 |  |  |
| 11 | 1.594E+21 | 1 |  |  |
| Jaccard distance | |  |  |  |
| Df | SumOfSqs | R^2^ | F | Pr(>F) |
| 3 | 1.24120418 | 0.72639605 | 7.07978139 | **0.0004** |
| 8 | 0.4675113 | 0.27360395 |  |  |
| 11 | 1.70871548 | 1 |  |  |

**Table S6. PERMANOVA results between Ciprofloxacin (CP) and control group (NC).**

| PhyloFunc distance | |  |  |  |
| --- | --- | --- | --- | --- |
| Df | SumOfSqs | R2 | F | Pr(>F) |
| 3 | 0.29579173 | **0.47158781** | **2.37989871** | 0.0148 |
| 8 | 0.33143341 | 0.52841219 |  |  |
| 11 | 0.62722513 | 1 |  |  |
| Bray-Curtis dissimilarity | |  |  |  |
| Df | SumOfSqs | R2 | F | Pr(>F) |
| 3 | 0.21199135 | 0.4435263 | 2.12541367 | 0.031 |
| 8 | 0.26597658 | 0.5564737 |  |  |
| 11 | 0.47796792 | 1 |  |  |
| Euclidean distance | |  |  |  |
| Df | SumOfSqs | R2 | F | Pr(>F) |
| 3 | 3.4339E+20 | 0.37451419 | 1.5966861 | 0.1479 |
| 8 | 5.7351E+20 | 0.62548581 |  |  |
| 11 | 9.169E+20 | 1 |  |  |
| Jaccard distance | |  |  |  |
| Df | SumOfSqs | R2 | F | Pr(>F) |
| 3 | 0.18612892 | 0.41984968 | 1.92984317 | **0.0027** |
| 8 | 0.25719384 | 0.58015032 |  |  |
| 11 | 0.44332276 | 1 |  |  |

**Table S7. PERMANOVA results between Azathioprine (AZ) and control group (NC).**
